# Natural genetic diversity in the potato resistance gene *RB* confers suppression avoidance from Phytophthora effector IPI-O4

**DOI:** 10.1101/2020.11.13.381905

**Authors:** Hari S. Karki, Sidrat Abdullah, Yu Chen, Dennis A. Halterman

## Abstract

*RB* is a potato gene that provides resistance to a broad spectrum of genotypes of the late blight pathogen *Phytophthora infestans*. *RB* belongs to the CC-NB-LRR (coiled-coil, nucleotide-binding, leucine-rich repeat) class of resistance (R) genes, a major component of the plant immune system. It directly interacts with Class I and II IPI-O effectors from *P. infestans* to initiate a hypersensitive resistance response, but this activity is suppressed in the presence of the Class III effector IPI-O4. Using natural genetic variation of RB within potato wild relatives, we identified two amino acids in the CC domain that alter interactions needed for suppression of resistance by IPI-O4. We have found that separate modification of these amino acids in RB can diminish or enhance the resistance capability of this protein against *P. infestans* in both *Nicotiana benthamiana* and potato. Our results demonstrate that increased knowledge of the molecular mechanisms that determine resistance activation and R protein suppression by effectors can be utilized to tailor-engineer genes with the potential to provide increased durability.

---

The majority of plant R proteins have nucleotide-binding (NB) and leucine-rich repeat (LRR) motifs and can be divided into two sub-classes based on their amino-terminal domain (Jones and Dangl, 2006). One subclass encodes a coil-coil (CC) domain, while the other contains homology to the Toll-interleukin-1 receptor (TIR) (Jones and Dangl, 2006). The N-terminal domain (TIR/CC) of plant NB-LRR proteins has been proposed to initiate signaling or recognition (Eitas and Dangl, 2010). Some host accessory proteins, which interact with both the R protein through the N-terminal domain and the cognate pathogen effectors, have been demonstrated to be required for disease resistance (Eitas and Dangl, 2010). To date, many effectors have been identified that can be recognized by the cognate host R proteins and activate effector-triggered immunity (ETI) upon recognition (Jones and Dangl, 2006). Foundational progress has also been made in elucidating the mechanisms of R protein/effector interaction. The “Guard Model”, “Decoy Model” and “Bait-and-Switch Model” have been proposed to explain the recognition mechanisms, and each of them are supported in multiple pathosystems (reviewed in van der Hoorn and Kamoun, 2008; Collier and Moffett, 2009; Paulus and van der Hoorn, 2018).

Late blight of potato, caused by the oomycete pathogen *Phytophthora infestans*, continues to be one of the most challenging diseases to sustainably and proactively manage. The most frequently used management strategy to control late blight is to spray fungicides repeatedly, which is expensive and causes environmental concerns. Therefore, the best way to control the disease is through natural host resistance. Wild potato species have provided a rich source of resistance against late blight. *Solanum demissum* is a hexaploid Mexican species from which 11 *R* genes have been identified, or cloned, and extensively used in potato breeding for late blight resistance. However, virulent races of *P. infestans* have rapidly overcome a majority of the *R* genes from *S. demissum* in most potato growing regions (Wastie, 1991). Three major late blight resistance genes have also been cloned from another diploid wild potato species *Solanum bulbocastanum*, including *RB* (Song et al., 2003), also known as *Rpi-blb1* (van der Vossen et al., 2003), *Rpi-blb2* (van der Vossen et al., 2005) and *Rpi-blb3* (Lokossou et al., 2009).

The *P. infestans* effector *Avrblb1*, also known as IPI-O (in planta induced gene O), was originally found from a screen for differentially expressed genes in *P. infestans* during potato infection (Pieterse et al., 1994; van West et al., 1998). IPI-O showed a high level of expression in invading hyphae (van West et al., 1998). Later studies revealed that IPI-O is a multigene effector family and the IPI-O locus can be extremely variable between isolates (van West et al., 1998; Champouret et al., 2009; Halterman et al., 2010). IPI-O variants have been divided into three classes based on diversity of their deduced amino acid sequences(Champouret et al., 2009; Halterman et al., 2010). In addition, *P. infestans* isolates lacking class I and class II variants are virulent on plants with *RB* (Champouret et al., 2009), and the *P. infestans* isolates with class III variants are more aggressive on plants with *RB* (Halterman et al., 2010). Interestingly, the class III variant IPI-O4 not only eludes detection by *RB*, but is also capable of inhibiting the HR elicited by class I variant IPI-O1 (Chen et al., 2012).

Despite the fact that *P. infestans* genotypes that overcome *RB* resistance have been found (Champouret et al., 2009; Halterman et al., 2010; Karki et al., 2020), *RB* remains valuable for potato breeding due to the almost ubiquitous existence of IPI-O1 in strains from major potato growing regions. We have previously shown that *RB*-containing plants respond through elicitation of an HR after inoculation with *P. infestans*, but differ in the transcription patterns for the HR-related gene *Hin1* and the PR genes *PR-1b*, *PR-2a*, and *PR-5* (Chen and Halterman, 2011). These observations explain to some extent the partial resistance phenotype that has been repeatedly observed in plants with the *RB* gene, and also suggest that *P. infestans* might be able to suppress responses in partially resistant plants with *RB* (Chen and Halterman, 2011). Together with our previous discovery that IPI-O4 is able to inhibit the HR elicited by IPI-O1 through directly binding the RB CC domain (Chen et al., 2012), the data indicate a complex and dynamic interface between potato *R* gene products and *P. infestans* effectors. In order to elucidate the molecular mechanisms underlying resistance suppression, we investigated the intra- and intermolecular interactions between the RB protein and IPI-O effectors and proposed a model based on our observations (Chen et al., 2012). The RB protein forms a multimeric complex through intermolecular interactions of the CC domain which is consistent with the ‘resistosome’ model proposed by (Wang et al., 2019a; Wang et al., 2019b). In the absence of IPI-O1, RB remains in a resting state. Upon recognition of IPI-O1, a conformational change occurs, which enables RB to activate resistance, possibly by exposing a platform for signaling components. However, when IPI-O4 is present, this effector interacts with the CC domain and disrupts CC oligomerization, or blocks an interaction with other signaling components, thus suppressing RB activation. This model illustrates that modification of the CC domain might be a useful strategy to engineer *RB* so that it can recognize more variants of IPI-O effectors, or alter intermolecular CC domain interactions to prevent disruption by IPI-O4.

The current work is focused on identifying variation within the RB CC domain that avoids interaction with IPI-O4 and thus could enable resistance activation even in the presence of this suppressor. We found that modification of single amino acids within the RB CC domain can either diminish the resistance capability of the modified RB or increase the ability of RB to protect plants against *P. infestans* lineages that have been shown to overcome the wild-type RB protein. Our results demonstrate that increased knowledge of the molecular mechanisms that determine resistance activation and R protein suppression by effectors can be utilized to tailor-engineer genes with the potential to provide increased durability.

## RESULTS

### RB CC domain amino acids 115 and 117 influence multimerization in yeast

We previously used natural variation of RB from wild species of potato to identify an RB CC domain form S. *pinnatisectum* (pnt) that does not interact directly with IPI-O1 or IPI-O4 in a yeast two-hybrid assay (Chen et al., 2012). Amino acid sequence alignment of the CC domains of wild type RB and the RB ortholog from *S. pinnatisectum* (pnt) identified 21 amino acids that were different between the two (Figure 1). Nine of these amino acids were non-conservative changes and were chosen for modification to test for their effect on intermolecular interactions. We performed site-directed mutagenesis to change the amino acids in the wild-type RB CC domain to the corresponding amino acids in the pnt RB. The modified RB CC domains were then tested for their ability to interact with the wild-type RB CC protein, IPI-O4, or self-interaction using a yeast two-hybrid assay (Table S1). We found that single amino acid changes K115T and K117D diminished the ability of the CC domain to interact with IPI-O4 (Table 1). These mutations also affected self-association of the CC domains and their ability to interact with the wild-type RB CC domain. Mutation K131N also diminished interaction between the modified and wild-type RB CC domains. The double mutant K115T/K117D also was not able to interact with IPI-O4 in yeast.

**Table 1.** Results of a yeast two-hybrid assay to test interactions of RB CC domain mutants with wild-type RB CC domain, itself, or IPI-O4.

| Amino acid mutation | RBCCblb | self | IPI-O4 |
| --- | --- | --- | --- |
| D29N | + | + | + |
| A43S | + | + | + |
| N55D | + | + | + |
| R85L | + | + | + |
| E90A | + | + | + |
| K115T | +/- | +/- | +/- |
| K117D | - | - | - |
| K131N | +/- | + | + |
| D155G | + | + | + |
| K158E | + | + | + |
| K115T, K117D | - | - | - |
| RBblb-CC wild-type | + | + | + |

**Figure 1.**
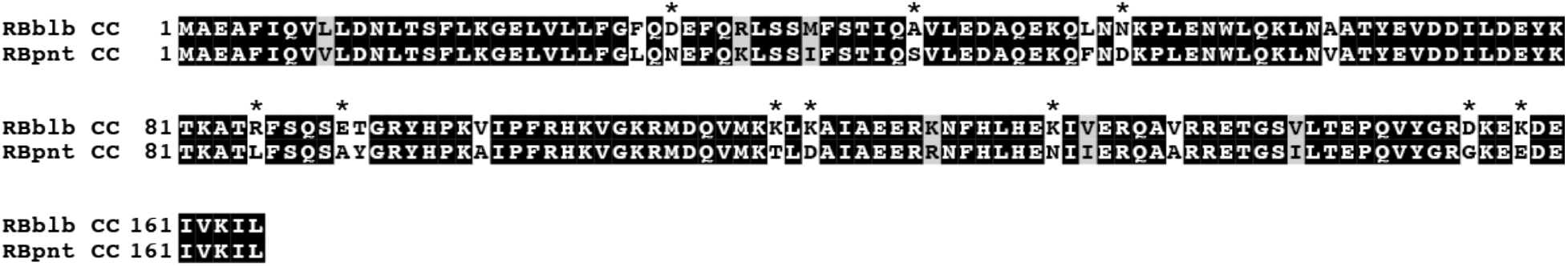
Amino acid alignment of the RB CC domain with the CC domain of the RB homolog from *S. pinnatisectum* (pnt). Amino acids highlighted in black, gray, and white indicate identical, similar, and non-conserved amino acids, respectively. Asterisks denote amino acids that were targeted for site-directed mutagenesis.

### RB CC domain amino acids 115 and 117 influence resistance activation

We tested whether changes in the protein at positions 115 and/or 117 within the RB CC domain still allow the protein to activate an HR elicited by IPI-O1. We created site-directed mutants at positions 115 and 117 of the wild type RB gene under the control of a native promoter and terminator. *N. benthamiana* leaves were co-agroinfiltrated with either *RB115*, *RB117*, or *RB115/RB117* and IPI-O1. The *RB117* mutant retained HR elucidating capability similar to the wild-type RB (Figure 2A). However, *RB115* completely lost its ability to elicit HR upon recognition of IPI-O1. Co-infiltration of the *RB115/RB117* double mutant with IPI-O1 also elicited an HR, but initiation of HR was delayed compared to wild-type *RB* and the *RB117* mutant. None of the *RB* genes induced an HR when co-infiltrated with IPI-O4 alone (Figure 2B), which is consistent with prior results and shows that there is no gain of IPI-O4 recognition conferred by the K115T or K117D mutations. Previously, we reported that IPI-O4 suppresses the HR upon the recognition of IPI-O1 by RB (Chen, 2012). In order to test whether RB117 escapes the suppression by IPI-O4, we co-infiltrated *N. benthamiana* leaves with either *RB*, *RB115*, *RB117*, or *RB115/RB117* and *IPI-O4*, followed by infiltration of *IPI-O1* at the same spot 24 hours later. Our results show that the *RB117* mutant retained its ability to elicit the HR upon recognition of IPI-O1 even in the presence of IPI-O4 (Figure 2C). Despite the ability of the double mutant to elicit an HR upon recognition of IPIO-1, we found that co-infiltration with *IPI-O4* led to suppression of the HR, suggesting that the 115 and 117 amino acids introduce subtle structural changes that modify the resistance protein’s interactions with these effectors.

**Figure 2.**
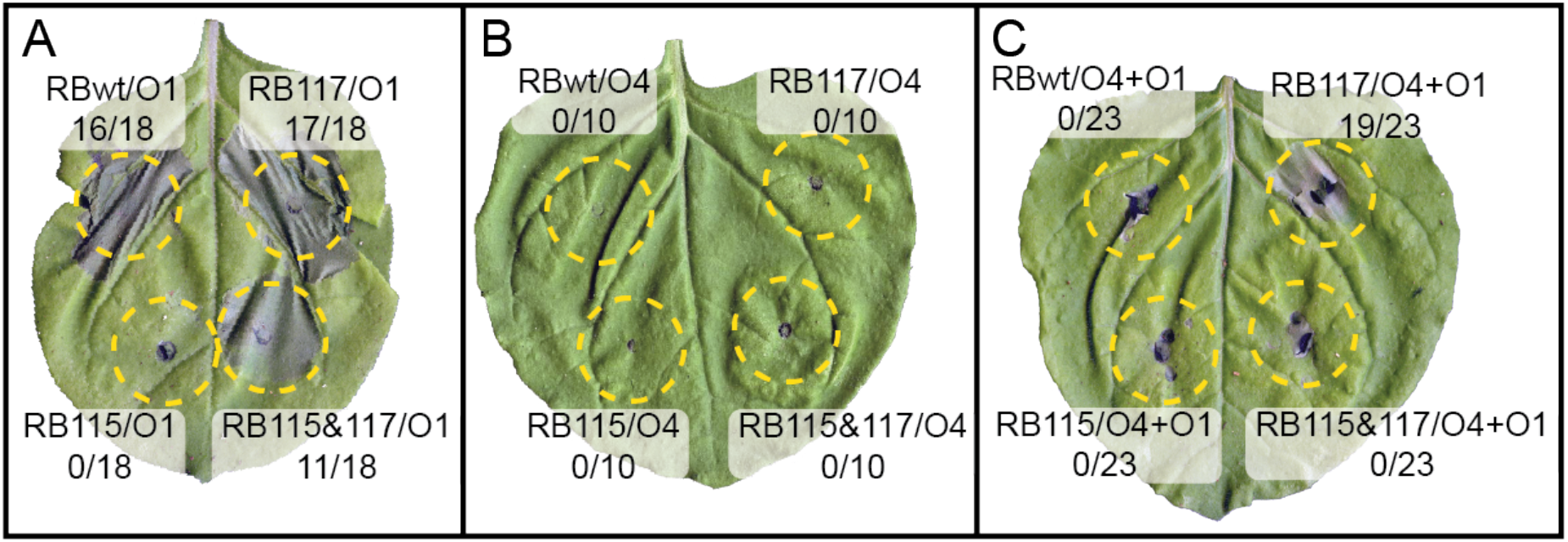
Effect of RB115 and RB117 mutations to elicit the HR upon recognition of *P. infestans* effectors IPI-O1 and IPI-O4. **A)** Wild-type RB (wt), RB115, RB117, and double mutant RB115/117 co-expressed with IPI-O1. **B)** RB (wild type), RB115, RB117 and double mutant RB115/117 co-expressed with IPI-O4. **C)** RB (wild type), RB115, RB117, and double mutant RB115/117 co-expressed with IPI-O4, followed by infiltration in the same spot with IPI-O1 after 24 hours. The numbers below each label indicate the number of attempts that exhibited cell death out of the total number of replications. Photographs were taken 6 days after the first infiltration.

### Introduction of the RB117 mutation does not affect resistance to *P. infestans* in *N. benthamiana*

To determine whether mutations at locations 115 and 117 in the RB protein affect resistance against *P. infestans*, we agroinfiltrated the altered genes into leaves of 4-5 week-old *N*. *benthamiana* followed by inoculation with *P. infestans* US-23. The modified *RB117* and *RB115/RB117* genes conferred resistance to *P. infestans*, compared to *RB115*, which was susceptible (Figure 3). The *RB117* gene provided slightly improved resistance compared to the wild-type *RB*, but this difference was not substantial. These constructs were also co-agroinfiltrated with the *IPI-O4* effector and tested for their ability to confer resistance to *P. infestans* US-23. Our results were consistent with our prior results showing that while IPI-O4 is capable of suppressing the elicitation of the HR, it is not able to block resistance to *P. infestans* completely (Chen and Halterman, 2017b). Regions that were co-agroinfiltrated with wild-type *RB*, *RB117*, or *RB115/117* along with *IPI-O4* were resistant, while co-agroinfiltration of *RB115* with *IPIO-4* remained susceptible (Figure 3).

**Figure 3.**
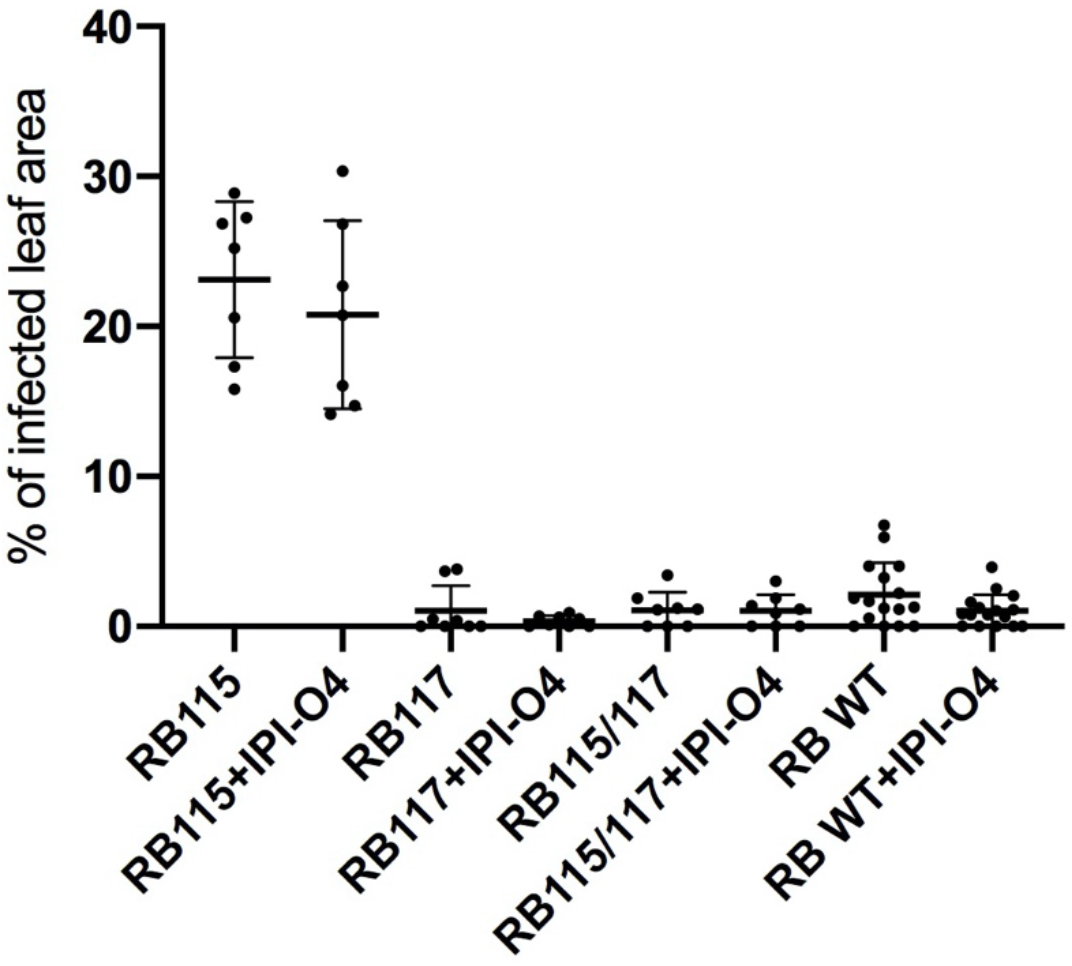
*P. infestans* resistance assay following transient expression of RB and RB mutants. Wild type *N. benthamiana* leaves were infiltrated with *A. tumefaciens* containing the following constructs: RB115, RB117, or RB wild type either alone or co-infiltrated with IPI-O4. At 24 hours after infiltration, the leaves were subjected to a detached leaf assay using *P. infestans* US-23. The percentage of infected leaf area was calculated 6 days after inoculation using ImageJ.

### *RB117* provides resistance in potato

Our results showing that *RB* with the K117D mutation confers resistance in *N. benthamiana* led us to test whether this gene is also functional against late blight in a potato pathosystem. We transformed potato cultivar Katahdin with either *RB115* or *RB117* under the control of an *RB*-native promoter and terminator. Although *RB115* was not able to confer resistance in the transient expression assay, we transferred it into potato to serve as a control in these experiments. We obtained 25 plantlets transformed with *RB115* and 27 plantlets transformed with *RB117* that successfully rooted on kanamycin-selective media. The plants were grown in the greenhouse and observed for true-to-type growth habits compared to non-transformed Katahdin. Of the 25 *RB115* transformants, ten survived in the greenhouse and exhibited a normal phenotype. Of the 27 *RB117* transformants, 25 had a true-to-type phenotype. Fully developed, young and healthy leaves from these plants were excised and used in a detached leaf assay with *P. infestans* isolate US-23. After 5 days, the inoculated leaves were scored for late blight symptoms (Figure 4). The majority of the RB115 plants were not significantly different from the susceptible wild-type Katahdin control. Three plants RB115(18), RB115(19), and RB115(20) all exhibited a resistance phenotype similar to the SP951 resistant control. However, repeat experiments using older leaves from these three RB115 individuals showed that they are susceptible to *P. infestans* (Figure S1), which is consistent with the previous assay using *N. benthamiana*. The majority of RB117 plants exhibited resistance although RB117(29), RB117(32), and RB117(39) showed the fewest symptoms. Repeat experiments using older leaves of these plants showed that they maintain resistance in older leaves (Figure S1).

**Figure 4.**
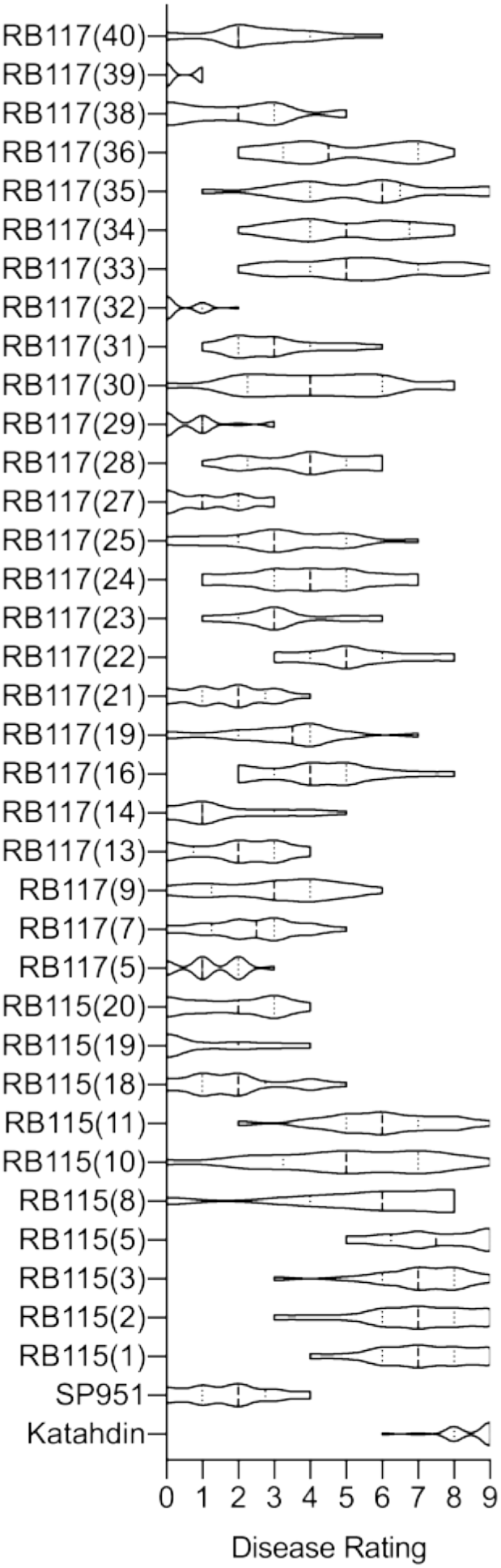
Results of a detached leaf assay using primary transformants of RB115 or RB117 in potato cultivar Katahdin. Numbers in parentheses denote the individual plantlet that was excised from transformed tissue. Disease rating data is represented as a violin graph. Dashed lines indicate the median and dotted lines denote the quartiles. The thickness of each plot represents the frequency distribution of the data.

### RB117 maintains resistance in potato despite the presence of IPI-O4

We previously showed that RB117 maintains its ability to elicit the HR and confer resistance to *P. infestans* in the presence of the IPI-O4 suppressor during transient co-expression in *N. benthamiana*. In order to test the effectiveness of RB117 in the presence of IPI-O4 in potato, we crossed the RB115(20) and RB117(39) plants with potato that is expressing the *IPI-O4* effector under the control of a 35S promoter (Chen and Halterman, 2017). Progeny that were PCR-positive for both *IPI-O4* and *RB* were used in a DLA to test for resistance to *P. infestans* US-23. Five progenies from the cross using RB115(20) and four progenies from the cross using RB117(39) were determined to contain *IPI-O4*. However, only one plant from each cross was found to have both *IPI-O4* and *RB*. These plants were used in a DLA to determine the resistance phenotype using *P. infestans* US-23 (Figure 5). The control plants, including susceptible Katahdin and Katahdin expressing IPI-O4, were both susceptible to *P. infestans*, while SP951 expressing the wild-type *RB* gene was considered resistant. Both RB115(20) and RB115(20)+IPI-O4 plants were also found to be susceptible. Consistent with our previous results, RB117(39) was resistant and showed fewer disease than SP951. The RB117(39)+IPI-O4 plant was also resistant and was not significantly different from the RB117(39) parent plant.

**Figure 5.**
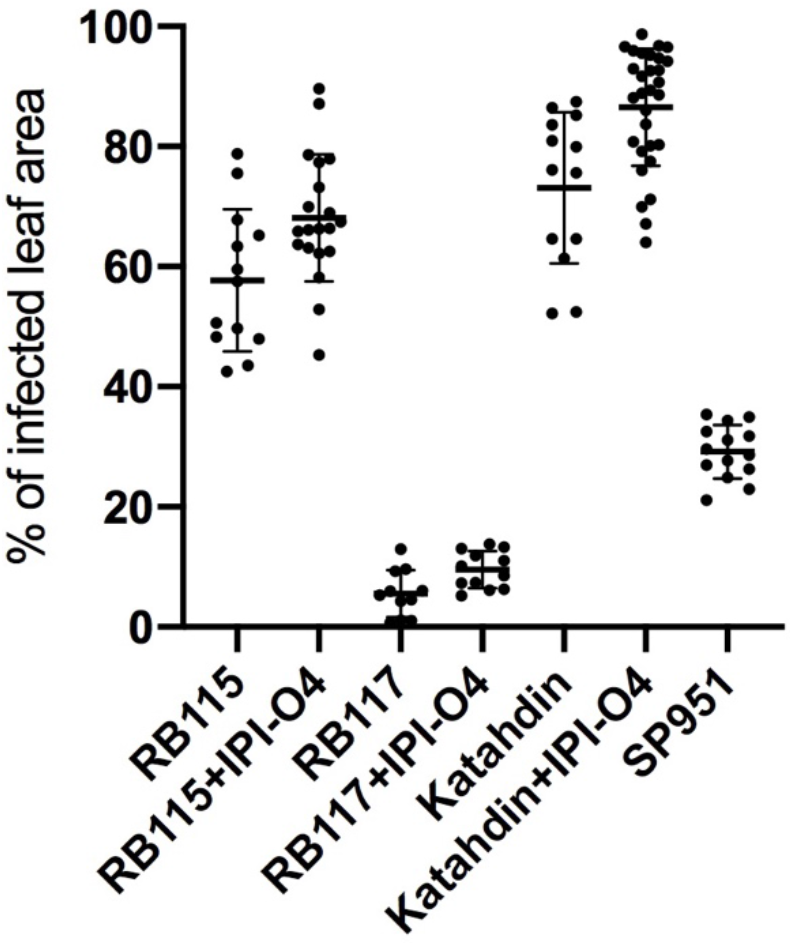
Detached leaf assay of potato leaves with *P. infestans* isolate US-23. Detached leaves were inoculated with *P. infestans* isolate US-23 by dropping 10 μl of a zoospore suspension on the abaxial side of the leaflets. Photographs were taken 5 days after inoculation and percentage diseased area of the leaflet was calculated using ImageJ.

### *RB117* does not provide resistance against *P. infestans* isolate NL13316 in potato

We hypothesized that the inability of IPI-O4 to suppress RB117 could lead to increased resistance to more aggressive *P. infestans* genotypes. To test this, we conducted a DLA in *N. benthamiana* and transgenic potato expressing *RB117* against the highly aggressive *P. infestans* isolate NL13316. This genotype has been shown to overcome RB resistance in potato (Karki et al., 2020). We first examined the susceptibility of *N. benthamiana* plants that had been stably transformed with the wild-type *RB* gene. We found that, similar to *RB* in potato, transgenic *N. benthamiana* was resistant to US-23, but susceptible to NL13316 (Figure 6A). We then carried out transient expression of wild-type *RB*, *RB115*, and *RB117* in *N. benthamiana* with and without IPI-O4, followed by inoculation with NL13316. We were surprised to find that transient expression of wild-type *RB* conferred resistance to NL13316, even in the presence of *IPI-O4* (Figure 6B). However, both *RB115* and *RB115*+*IPI-O4* were susceptible to NL13316, ruling out the possibility of agrobacterium influencing the outcome of the interactions. Areas infiltrated with *RB117* or the double mutant, with or without *IPI-O4*, were all resistant to NL13316 at a level similar to the wild-type *RB*.

**Figure 6.**
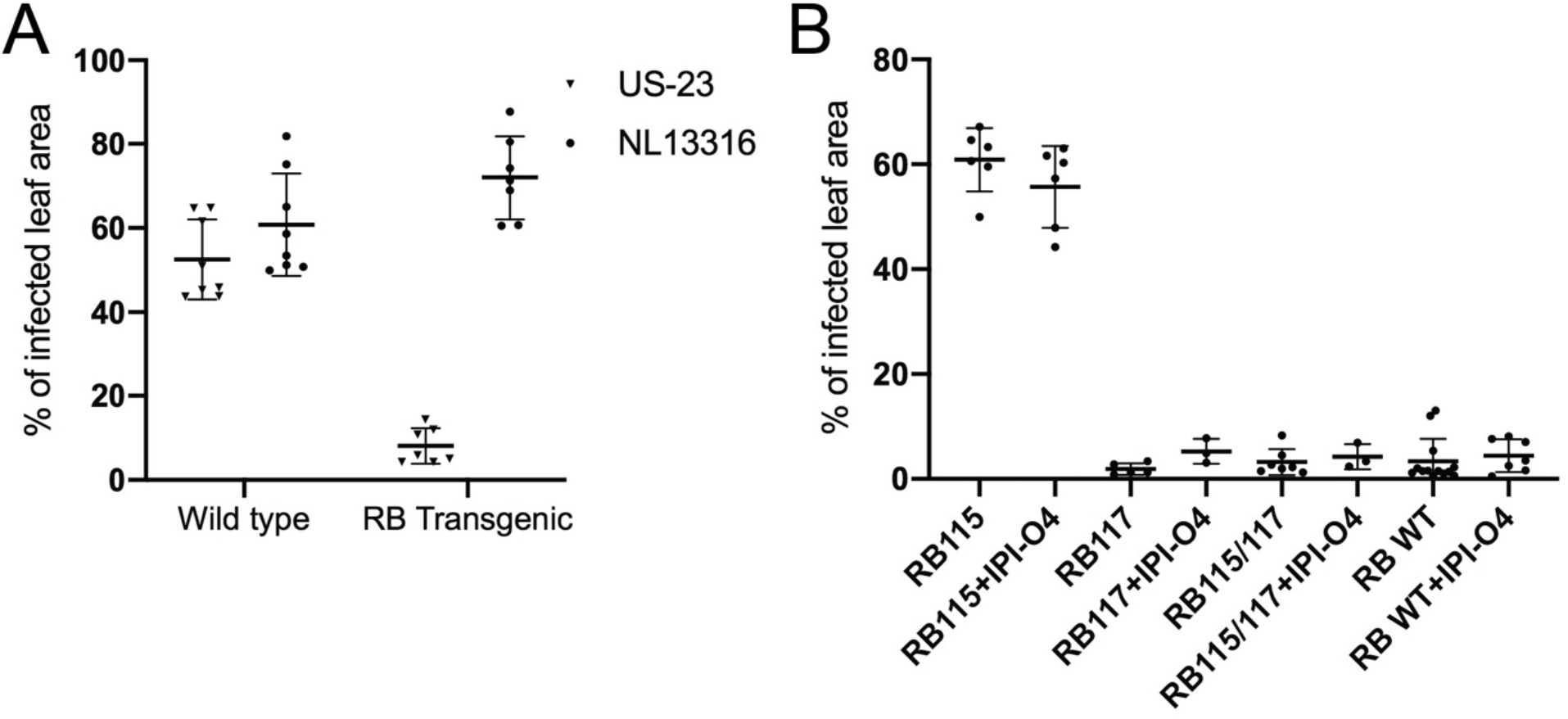
Disease assay on non-transgenic (Wild type) and RB transgenic *N. benthamiana*. **A)** Fully developed 4-week-old leaves were inoculated with the *P. infestans* isolates US-23 and NL13316. The percentage of the infected leaf area was calculated 5 days after infection using ImageJ. **B)** Wild type *N. benthamiana* leaves were infiltrated with *A. tumefaciens* containing either RB115, RB117, or RB WT with or without co-infiltration with IPI-O4. At 24 hrs after infiltration, the leaves were collected and subjected to a detached leaf assay using *P. infestans* isolate NL13316. The percentage of the infected leaf area was calculated 6 days after infection.

In potato, a DLA was conducted using Katahdin+*RB117*, Katahdin+*RB115*, SP951, K41 (expressing the wild-type RB gene derived from somatic fusion with *S. bulbocastanum*), and non-transformed Katahdin inoculated with US-23 and NL13316 side by side on the same leaves (Figure 7). Following inoculation with *P. infestans* US-23, the *RB117* and K41 leaves showed resistance, and SP951 showed partial resistance, while *RB115* and wild-type Katahdin were susceptible. We observed that along with K41 and SP951, *RB117* was also susceptible to NL13316. These results indicate that although *RB117* is able to escape *IPIO-4* suppression and provide adequate resistance to US-23, this gene does not provide resistance against *P. infestans* isolate NL13316 in potato.

**Figure 7.**
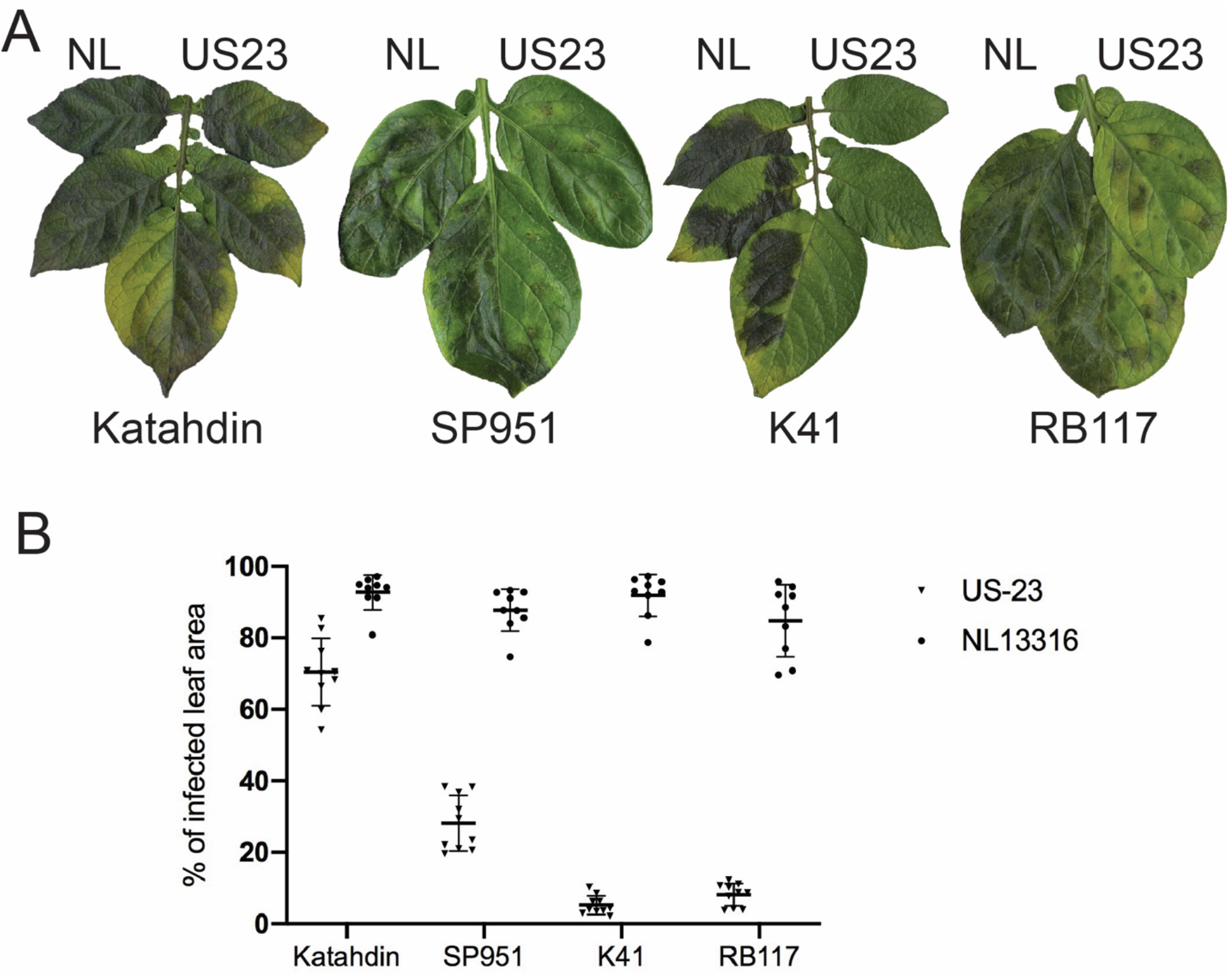
Detached leaf assay of potatoes with *P. infestans* US-23 and NL13316. Each leaf was inoculated with US-23 on the left and NL13316 (NL) on the right. Photographs were taken 5 days after inoculation and the percentage of diseased area of the leaflet was calculated using ImageJ. **A)** Representative photos of leaves from the detached leaf assay. **B)** Percentages of the diseased leaf areas.

## DISCUSSION

To suppress PAMP-triggered immunity (PTI), pathogens introduce effector proteins into and around host cells (Stergiopoulos and Wit, 2009; de Jonge et al., 2011; Giraldo and Valent, 2013). Cytoplasmic pathogen effectors are recognized by intracellular nucleotide-binding leucine-rich repeat (NLR) receptors that trigger effector-triggered immunity (ETI; Jones and Dangl, 2006; Dodds and Rathjen, 2010). Effector recognition by R proteins is critical for the determination of resistance and susceptible phenotypes in plants by the pathogens. In this study, we have identified amino acids in the CC domain of the RB protein that are critical for molecular interactions that determine effector recognition and suppression by the *P. infestans* effector IPI-O4.

RB recognizes IPI-O variants belonging to class I (IPI-O1, IPI-O2, IPI-O5, IPI-O7, and IPI-O8) and class II (Champouret et al., 2009) to initiate a hypersensitive response (HR), but does not recognize class III (IPI-O4; Champouret et al., 2009; Halterman et al., 2010). Instead, IPI-O4 blocks RB-mediated recognition of IPI-O1 to induce cell death in *N. benthamiana* and IPI-O4 expressing *P. infestans* can overcome *RB* mediated resistance in potato (Halterman et al., 2010). Like Tobacco N, Arabidopsis RPP1 and Flax L5, L6, and L7 proteins (Erickson et al., 1999; Dodds et al., 2006; Krasileva et al., 2010), RB directly interacts with the IPI-O1 effector (Chen et al., 2012). The RB CC domain undergoes self-association and also physically interacts with IPI-O1 and IPI-O4 to turn on/off resistance (Chen et al., 2012). The role of the CC domain in the function of CC-NB-LRR proteins varies greatly. For example, the CC domain of RPM1 interacts with a host accessory protein RIN4 in Arabidopsis, which physically interacts with the cognate effectors *AvrRpm1* and *AvrB* (Mackey et al., 2002). In tomato, a CC-NB-LRR type of R protein Prf interacts with a host accessory protein Pto through the CC domain, and Pto interacts with the cognate effectors AvrPto or AvrPtoB physically (Mucyn et al., 2006). Overexpression of the CC domain of barley MLA can induce cell death in *N. benthamiana* and interact with WRKY transcription factors to inhibit their ability to repress defense genes (Shen et al., 2007). Similarly, the CC domain of NRG1- and ADR1-like proteins of Solanaceous species and *Arabidopsis thaliana* are sufficient to induce defense responses (Collier et al., 2011). In contrast, overexpression of the Rx CC domain alone does not induce an HR in *N. benthamiana*, however along with the LRR domain it co-regulates the signaling activity of the NB domain (Rairdan et al., 2008).

The CC domain of RB is essential for HR elicitation in *N. benthamiana* and potato through its interaction with IPI-O1 (Chen et al., 2012). We have previously shown that IPI-O effectors such as IPI-O1 and IPI-O4 are able to interact with the vast majority of CC domains of RB from wild potatoes (Chen et al., 2012). The wild species accessions that harbored these RB CC domains do not contain functional full-length RB homologs and were only used as a source of natural genetic variation. Since interaction between IPI-O4 and the RB CC domain induces suppression of resistance, we used protein-protein interaction to screen for variation that could potentially escape IPI-O4 suppression. The CC domain of an RB homolog from *S. pinnatisectum* (pnt), RBpnt, does not interact directly with IPI-O1 and IPI-O4 in yeast (Chen et al., 2012). Sequence variation between the CC domains of RB and RBpnt identified 21 amino acid differences, nine of which conferred non-conservative changes. Using site-directed mutagenesis, we found that two of these amino acids at locations 115 and 117 affect CC domain dimerization and interaction with IPI-O4. Furthermore, we showed that amino acid 117 is likely a target for IPI-O4 suppression, as modification of the RB CC domain at this location allowed for resistance activation even in the presence of IPI-O4. Comprehensive mutational analysis in the Arabidopsis *RPS2* and *Pseudomonas syringae avr-Rpt2* system has shown that single amino acid substitution in *RPS2* can disrupt cell death mediated by the R protein/effector interaction (Axtell et al., 2001). Our result is in the line with previous findings that targeted mutations in the R protein can expand the effector recognition spectrum (Farnham and Baulcombe, 2006; Harris et al., 2013; Segretin et al., 2014). Random mutagenesis of the potato NLR resistance protein R3a identified variation that allowed it to recognize the normally undetectable AVR3a^EM^ effector variant from *P. infestans* while still maintaining recognition of the AVR3a^KI^ isoform (Segretin et al., 2014). This mutation, located within the nucleotide-binding pocket, resulted in a gain-of-function response in transient assays with the effectors in *N. benthamiana*, but failed to improve resistance to *P. infestans* after transient or stable expression *N. benthamiana* or potato (Segretin et al., 2014). Here, we found that stable expression of RB117 is able to provide resistance to *P. infestans* US-23 after stable expression in potato but is unable to withstand the RB-breaking genotype NL13316. Random mutagenesis of the potato Rx NLR protein identified variation that provided broader-spectrum resistance to Potato Virus X but led to an increased fitness cost to the host (Farnham and Baulcombe, 2006). Further mutational analysis identified variation within the R protein that could offset this fitness cost and led to a stronger resistance response (Harris et al., 2013). Despite these apparent limitations, modified R proteins that potentially expand their recognition spectrum are still useful in crop improvement. The combination of wild-type and modified R proteins could lead to an optimized defense against pathogens and make it more difficult for the pathogen to evolve to avoid recognition or suppress resistance.

The *RB* gene provides broad-spectrum resistance to a range of *P. infestans* genotypes (Song et al., 2003; van der Vossen et al., 2003). However, there are multiple reports of *P. infestans* isolates that can overcome the *RB* gene (Champouret et al., 2009; Halterman et al., 2010; Karki et al., 2020). Strategies to combat resistance-breaking genotypes of *P. infestans* can include stacking of R genes that confer recognition of essential effectors, or effectors expressed at different developmental stages of the pathogen. Additionally, we provide support for an approach that includes improved R proteins that can avoid effector suppression or broaden recognition specificity. Our results indicate that although RB117 escapes IPI-O4 suppression, and is effective at limiting infection of *P. infestans* US-23, it did not provide adequate resistance against the *RB*-breaking genotype NL13316 in potato. The US-23 genome contains the IPI-O4 effector (Chen and Halterman, 2017a) even though the wild-type *RB* gene is still effective against this genotype. Additionally, even overexpression of IPI-O4 in potato does not completely disrupt resistance in plants that contain RB (Chen and Halterman, 2017b). This suggests that other factors within the pathogen can modulate the effectiveness of resistance suppression and virulence. The NL13316 genotype is relatively uncharacterized and we look forward to understanding more about its ability to overcome RB-mediated resistance. Interestingly, we found that NL13316 could not overcome transiently expressed wild-type RB or RB117 in *N. benthamiana*. This further suggests that multiple factors beyond the binary RB/IPI-O interaction are necessary to determine the outcome of the host-pathogen encounter.

Late blight still remains the devastating disease in the potato production system. The cloning of Rpi genes offers the possibility of using them in potato improvement programs using biotechnology. However, the lifespan of R genes is determined by the ability of the pathogen to evolve to avoid recognition or suppress defense responses. Molecular and mutational characterization of *RB* and its corresponding effector presented in this study demonstrate that engineering new alleles could strengthen the arsenal of R genes available for resistance breeding. Current advancements in genome editing technology can target specific nucleotide sequences and could provide an avenue to modify R genes without the requirement of the transferring non-plant sequences. Further functional and biochemical understanding of RB117 with several *P. infestans* isolates will assist in the use of this gene in late blight management and increased resistance durability.

## MATERIALS AND METHODS

### Plant materials and growth conditions

Potato plants were grown in the greenhouse in soil-less potting mix (BM1, Berger) under greenhouse conditions (23°C day/15°C night temperatures with 14 hours of light). Wild type and *RB*-transgenic *N. benthamiana* plants were grown in pots in a growth chamber with 28°C day/27°C night temperatures and 16 hours of light. All the plants were watered and fertilized regularly as needed.

### Yeast two-hybrid assay and site-directed mutagenesis

The CytoTrap® system (Stratagene) was used to detect protein-protein interactions. Constructs used in this paper to test interactions are the same as published previously (Chen et al., 2012). Site directed mutagenesis of RB CC domains for interaction testing was done using the wild-type RB CC domain in the pMyr vector as a template. A pair of oligonucleotide primers containing the desired nucleotide substitution, each complementary to the opposite strands of the same target sequence, were extended during temperature cycling by using *PfuTurbo* DNA polymerase (Stratagene). A list of designed primers is shown in Table S1. The PCR conditions for substitution mutants were: 2 min at 95°C followed by 18 cycles of 30-50 sec at 95°C, 50 sec at 72-75°C, 13 min at 68°C, and 7 min at 68°C. After the temperature cycling, the PCR products were treated with *DpnI* for 2 hr at 37°C to digest the methylated parental DNA template. The resulting products were transformed into GC5^TM^ chemical competent cells (GeneChoice®). Mutations at the targeted sites were verified by sequencing.

Mutations at locations 115 and 117 in the full-length RB gene were generated by circular PCR (as outlined above) using the RGA2-PCR long-range PCR product that was first cloned into the BamHI site of pBluescript KS. The resultant mutated plasmid constructs were verified by sequencing before subcloning the modified RB gene into the BamHI site of the pCLD04541 vector and introduction into *Agrobacterium tumefaciens* GV3101.

### Generation of transgenic potatoes

The RB modified versions, RB115 and RB117, were transferred into potato cultivar Katahdin as described previously (Bhaskar et al., 2008). The transgenic plants were maintained in tissue culture media and propagated by cuttings in the greenhouse. The introduction of these genes was confirmed using RB specific primers (Table 2) and PCR conditions were as follows: 1 min at 94°C followed by 35 cycles of 15 s at 94°C, 30 s at 55°C, 30 s at 68°C, and 7 min at 68°C. Genomic DNA was extracted from the young growing leaves of potato plants with the DNeasy Plant Maxi Kit (QIAGEN, Hilden, Germany) according to the manufacturer instructions. To transfer the RB117 into IPI-O4 genetic background (Chen and Halterman 2017 phytopathology), we crossed the selected RB117 and RB115 transgenic potatoes to the IPI-O4 containing transgenic potato. The resulting F1 progenies were tested for the presence of IPI-O4 using primers specific for the effector and for RB (Table 2).

### *P. infestans* isolates and late blight infection assay

*P. infestans* isolates US-23 and NL13316 were used for late blight infection in a detached leaf assay. US-23 and NL13316 were kindly provided by A. Gevens (University of Wisconsin, Madison) and Simplot Plant Sciences, respectively. *P. infestans* growth conditions, zoospore production and infection assays were performed as described previously (Karki et al., 2020). Infected area in the leaves were quantified using ImageJ software (Rueden et al., 2017).

### Disease and HR assay in *N. benthamiana*

A 5 ml culture of *A. tumefaciens* carrying the various constructs was grown overnight at 28°C in media supplemented with proper antibiotics. *Agrobacterium* cells were harvested and resuspended in MMA induction buffer (1 liter MMA: 1g MS salts, 1.95 g MES, 20g sucrose, 200 μM acetosyringone, pH 5.6). All suspensions were incubated at room temperature for 2 hr prior to infiltration. For the disease (late blight infection) assay, *A. tumefaciens* that had been transformed with the *R* genes at a concentration of OD_600_ = 0.3 were infiltrated in the *N. benthamiana* leaves. At 24 hours after infiltration a detached leaf assay was performed with *P. infestans* isolates US-23 and NL13316 with zoospore concentrations of 50,000 per ml. The leaves were photographed 6 days after inoculation unless otherwise stated, and infected areas in the leaves were quantified using ImageJ (Rueden et al., 2017).

For the assay, R genes and effectors were equally combined to a final OD_600_ of 0.3. For the HR suppression assay by IPI-O4, first RB, RB115, and RB117 and IPI-O4 were co-infiltrated then 24 hours later IPI-O1 was infiltrated on the same spot as in (Chen et al., 2012). The infiltrated leaves were assessed for cell death and were photographed after 6 days. Two leaves from three to four plants were tested for each treatment and the experiment was replicated three times.

## ACKNOWLEDGEMENTS AND DISCLAIMER

Funding was provided by USDA-NIFA Plant-Associated Microbes and Plant-Microbe Interactions Award Number 2014-67013-21593; USDA-NIFA/NSF Plant Biotic Interactions Program award number 2018-67014-28488; ARS-State Cooperative Potato Research Program. This research was supported in part by the U.S. Department of Agriculture, Agricultural Research Service. The findings and conclusions in this publication are those of the author(s) and should not be construed to represent any official USDA or U.S. Government determination or policy.

---

**Figure S1.**
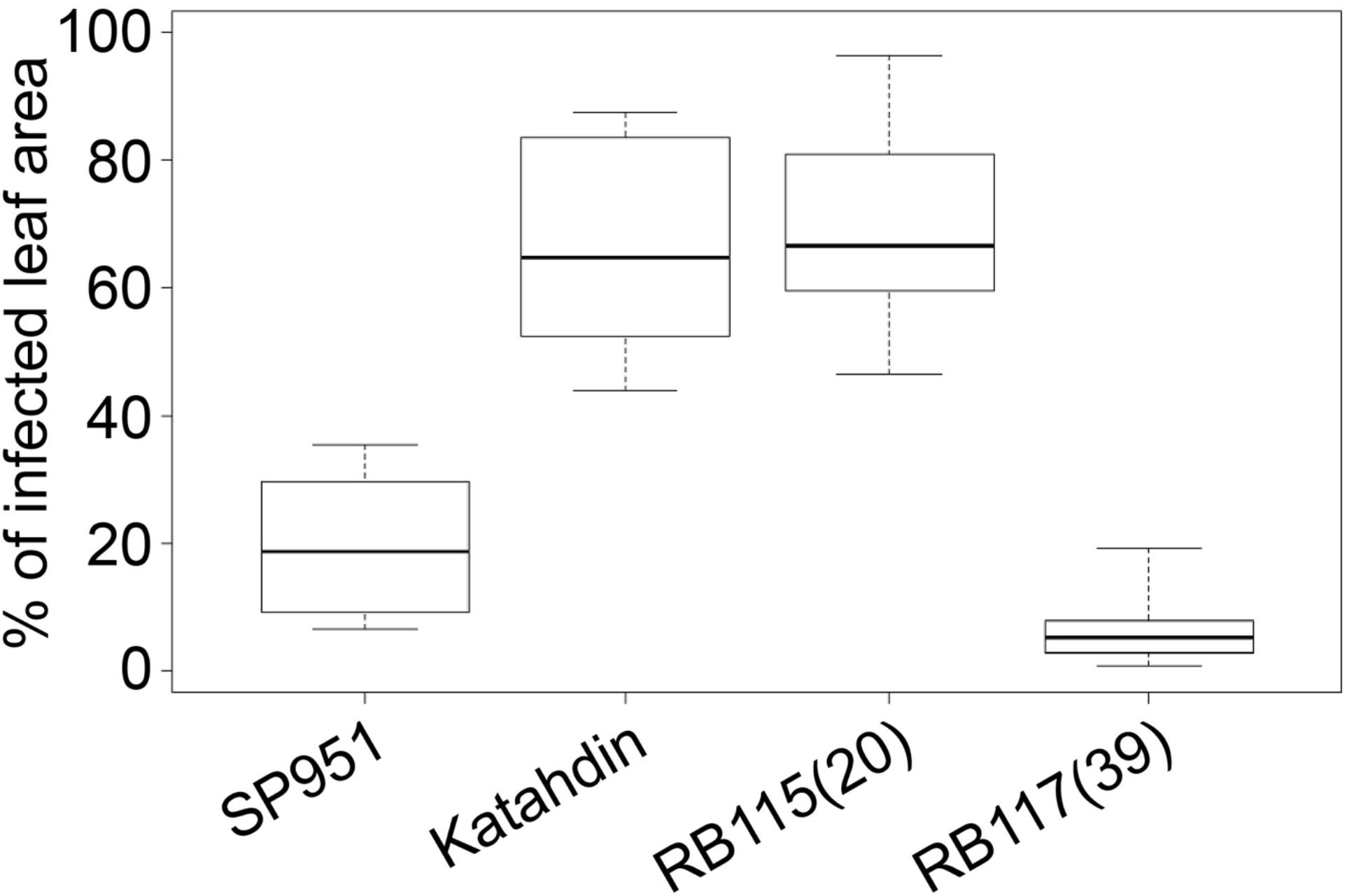
Late blight infection assay with *P. infestans* isolate US-23. Transgenic events harboring RB115 and RB117 gene were selected and used in a late blight infection assay. Photographs were taken 5 days after inoculation and percentage diseased area of the leaflet was calculated using ImageJ.

**Table S1.** Yeast two-hybrid results to test interactions between RB CC domain mutants with IPI-O and wild-type RB CC.

| Construct | AA mutation locations | IPIO-1 | IPI-O4 | wild-type RB CCblb |
| --- | --- | --- | --- | --- |
| 1 | 29, 43, 85, 90 | - | + | +/- |
| 2 | 43, 55, 85, 90 | - | + | + |
| 3 | 43, 85, 90, 117 | - | + | - |
| 4 | 43, 85, 90, 131 | - | + | + |
| 5 | 43, 85, 90, 158 | - | + | N/D |
| 6 | 29, 43, 85, 115 | - | + | - |
| 7 | 43, 55, 85, 115 | - | + | +/- |
| 8 | 43, 85, 117 | - | + | - |
| 9 | 43, 85, 115, 131 | - | + | - |
| 10 | 43, 85, 115, 158 | - | + | +/- |
| 11 | 29, 43, 55, 85, 90 | - | + | + |
| 12 | 29, 43, 85, 90, 115 | - | + | - |
| 13 | 29, 43, 85, 90, 117 | - | + | - |
| 14 | 29, 43, 85, 90, 131 | - | + | + |
| 15 | 29, 43, 85, 90, 158 | - | + | + |
| 16 | 29, 43, 55, 85, 90, 115 | - | + | - |
| 17 | 29, 43, 55, 85, 90, 117 | - | + | - |
| 18 | 29, 43, 55, 85, 90, 131 | - | + | + |
| 19 | 29, 43, 55, 85, 90, 158 | - | + | + |
| 20 | 29, 43, 85, 90, 115, 131 | - | + | - |
| 21 | 29, 43, 85, 90, 115, 158 | - | + | +/- |
| 22 | 29, 43, 85, 90, 117, 131 | - | + | - |
| 23 | 29, 43, 85, 90, 117, 158 | - | + | - |
| 24 | 29, 43, 85, 90, 131, 158 | - | + | + |
| 25 | 29, 43, 55, 85, 90, 115, 117, 131 | - | + | - |
| 26 | 29, 43, 55, 85, 90, 115, 117, 158 | - | + | - |
| 27 | 29, 43, 55, 85, 90, 115, 117, 131, 158 | - | + | - |
| 28 | RB-CCpnt wild-type | - | - | - |
| 29 | RB-CCblb wild-type | - | + | + |

**Table S2.**
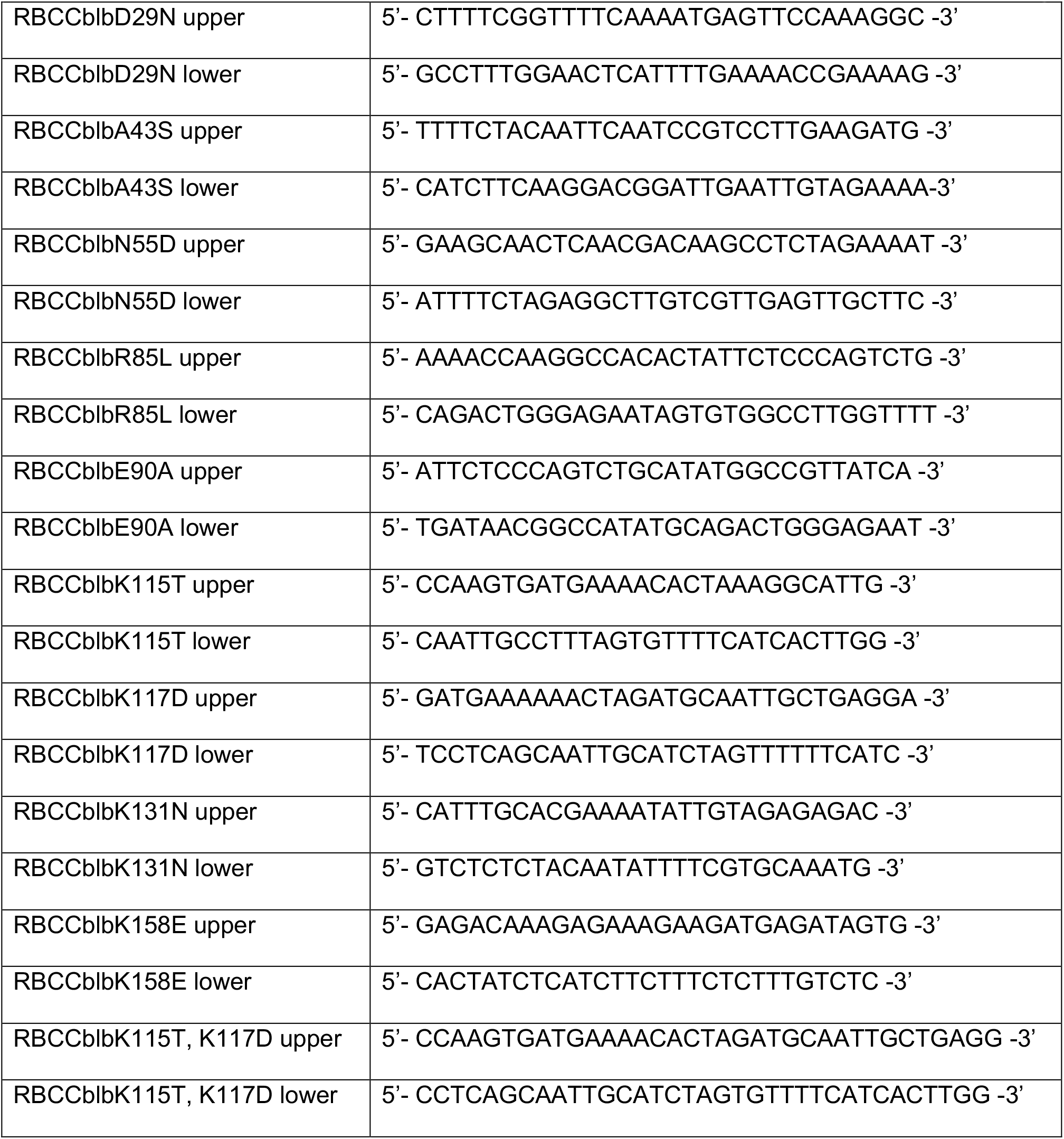
Primers used for site-directed mutagenesis of RB CC domain for yeast two-hybrid experiments.

